# Immune cell networks uncover candidate biomarkers of cancer immunotherapy response

**DOI:** 10.1101/2021.04.22.440902

**Authors:** Duong H.T. Vo, Gerard McGleave, Ian M. Overton

**Affiliations:** The Patrick G Johnston Centre for Cancer Research, Queen’s University Belfast, 97 Lisburn Road, BT9 7AE, United Kingdom

**Keywords:** Immune checkpoint, melanoma, ovarian carcinoma, systems immunology, network biology, immunotherapy, precision oncology, biomarker, nivolumab, systems medicine

## Abstract

Therapeutic activation of antitumour immunity by immune checkpoint inhibitors (ICIs) is a significant advance in cancer medicine, not least due to the prospect of long-term remission. However, many patients are unresponsive to ICI therapy and may experience serious side effects; companion biomarkers are urgently needed to help inform ICI prescribing decisions. We present the IMMUNETS networks of gene coregulation in five key immune cell types, and application to interrogate control of nivolumab response in advanced melanoma cohorts. Results evidence a role for each of the IMMUNETS cell types in ICI response and in driving tumour clearance with independent cohorts from TCGA. As expected, ‘immune hot’ status, including T cell proliferation, confers good response to first line ICI therapy. Genes regulated in NK, dendritic and B cells are the most prominent discriminators of nivolumab response in patients that had previously progressed on another ICI. Multivariate analysis controlling for tumour stage and age highlights B cell genes as candidate prognostic biomarkers. IMMUNETS provide a resource for network biology, enabling context-specific analysis of immune components in orthogonal datasets. Overall, our results illuminate the relationship between the tumour microenvironment and clinical trajectories, with potential implications for precision medicine.

## Introduction

The immune system functions to eliminate tumour cells; however, the immune response may also promote cancer progression, as a result of immunoediting [1]. The ‘equilibrium’ and subsequent ‘escape’ phases of immunoediting involve a selective microenvironment where cancer cells with capacity to evade the immune response become dominant. Therefore, tumour-associated leukocytes can adopt a range of different cellular programmes that may either impede or contribute to cancer progression [2]. Therapeutic activation of antitumour immunity has had huge clinical impact, particularly in producing long-term remission [3]. Immune checkpoint inhibitors (ICIs) exploit the CTLA-4 and PD-L/PD-L1 pathways to activate T lymphocytes in the tumor microenvironment [3]. ICIs are effective in a range of cancers, including melanoma, renal cell carcinoma and non-small cell lung cancer [4]. However, current forms of ICI therapy are also associated with multiple side effects, including autoimmune reactions [5] and response rate varies between cancer types, although a minority of patients typically respond to single-agent treatment [6]. Factors that correlate with ICI response include high mutational burden, a T cell-inflamed microenvironment and the formation of tertiary lymphoid structures with CD20^+^ B-cells [7,8]. Deeper understanding of the factors that drive the heterogeneity of clinical response to ICI therapy could inform patient-specific regimes in order to enhance outcomes and reduce side effects [9]. Indeed, treatment efficacy may be enhanced by combination therapy; for example, by targeting multiple pathways or by inhibition of tumour plasticity alongside ICIs [10,11]. An early successful example of a precision oncology approach measures HER2/neu in order to guide entry of breast cancer patients into the Trastuzumab treatment pathway [12]; similarly, development of companion biomarkers is a key step in maximising patient benefit from ICI therapy.

Immunotherapy has been particularly successful in melanoma, which is a common form of skin cancer associated with a high mortality rate due to its propensity to metastasise [13]. However, many melanoma patients do not benefit from sustained responses to ICIs, with median progression at 12 months or less [14–16]. Ovarian cancer also has poor prognosis, partly because around 70% of cases are diagnosed at an advanced stage [17]. Surgery and platinum-based chemotherapy are standard treatments for ovarian cancer, although 10-15% of patients respond to single-agent treatment with ICIs [18]. Characterisation of the molecular networks that control resistance and response to ICI therapy in poor prognosis cancers may provide mechanistic insight towards new therapeutic tools and ultimately inform prescribing decisions. Network models provide a useful abstraction of complex biological systems [19] and may enhance the development of methods for risk stratification in precision oncology [20]. Analysing data from the IRIS study [21] and melanoma patients treated with nivolumab, a PD-1 inhibitor [22,23], we map the activation and differentiation of five different immune cell types in order to identify molecular correlates of clinical immunotherapy responses. Our results delineate immunological processes at genome scale and propose candidate immunotherapy response biomarkers that have prognostic value in independent cohorts of cutaneous melanoma and ovarian carcinoma patients.

## Results

We produced IMMUNETS; a set of five immune cell coregulation networks derived from the Immune Response *In Silico* (IRIS) gene expression microarray data, representing cells from healthy human donors [21]. The IRIS study captured immune transcriptomes across multiple activated and differentiated cell states and therefore offers an excellent starting point for modelling the networks of genes involved in immune regulation and immunotherapy response. Our approach (Figure 1) utilized transcriptome data from IRIS, from melanoma patients treated with immunotherapy [22] and from The Cancer Genome Atlas (TCGA) [24,25]. IMMUNETS provided a basis for derivation of immune cell-specific focus networks (Figure 2, Supplementary Figures S1, S2, S3) and investigation of immune regulation in the context of nivolumab response (Figure 3). The prognostic value of selected candidate biomarkers was confirmed in the independent TCGA melanoma and ovarian cancer datasets (Tables 3, 4, 5). Ovarian cancer was selected as an additional comparator for survival analysis because it is a poor prognosis cancer where immunotherapy has value [17, 18].

**Figure 1.**
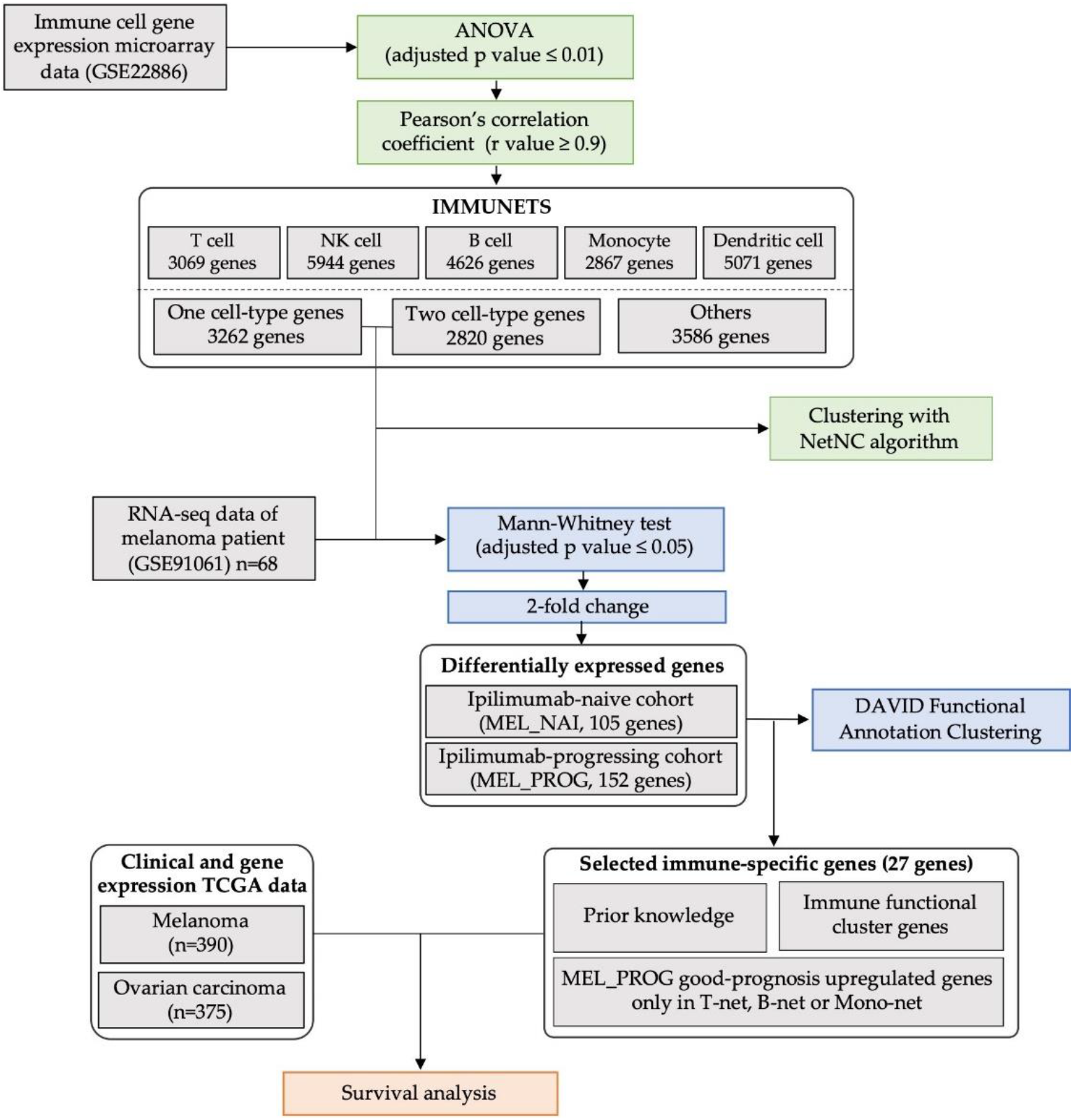
Methodological outline. Networks were constructed from genes that were both highly correlated and differentially expressed across five cell types (IMMUNETS) (centre, top). Genes present in at most two of the five IMMUNETS cell networks were input to the NetNC algorithm and were used to investigate immune regulation in patients with differing nivolumab response profiles (centre). IMMUNETS genes that correlated with nivolumab response were investigated in survival analysis with TGCA melanoma and ovarian carcinoma datasets (bottom).

**Figure 2.**
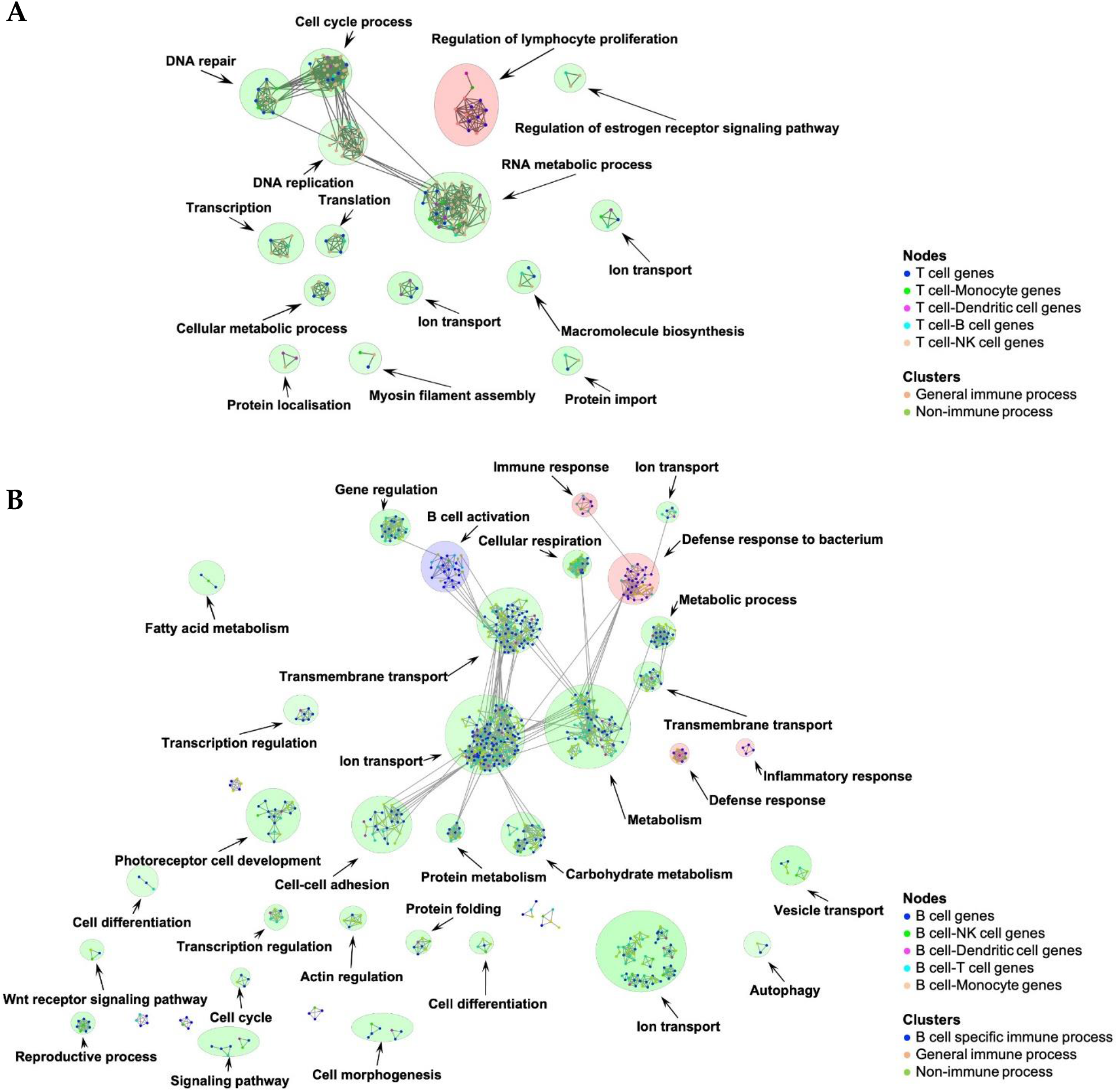
T cell and B cell focus networks. Node colour indicates the representation of genes in the IMMUNETS networks across the five different cell types examined, where blue shows genes present in a single network; the T cell and B cell networks are shown. Edges were output by NetNC-FTI (please see methods). The coloured circles around clusters indicate broad annotation classes; red corresponds to general immune GO annotations, blue represents cell-type specific immune processes and green shows clusters with annotation terms that are not immune-specific. The T cell focus network is shown in (A) where most clusters represent processes important for cell proliferation. A ‘regulation of lymphocyte proliferation’ cluster (red) contains multiple T cell genes (IL17A, CD5, IL13, IL5, IL2, CD8A, CD28). The B cell focus network is shown in (B) and has one cell-specific immune cluster (blue): ‘B cell activation’, including CD79A, CD79B and CD19. The four clusters annotated with general immune processes (red), such as ‘inflammatory response’, contain important B cell genes including IL22, IL10RB, IL20RA and IL22RA1. Clusters with significant GO terms are shown. Cytoscape sessions for the IMMUNETS focus networks are available in Supplementary Data File S2.

**Figure 3.**
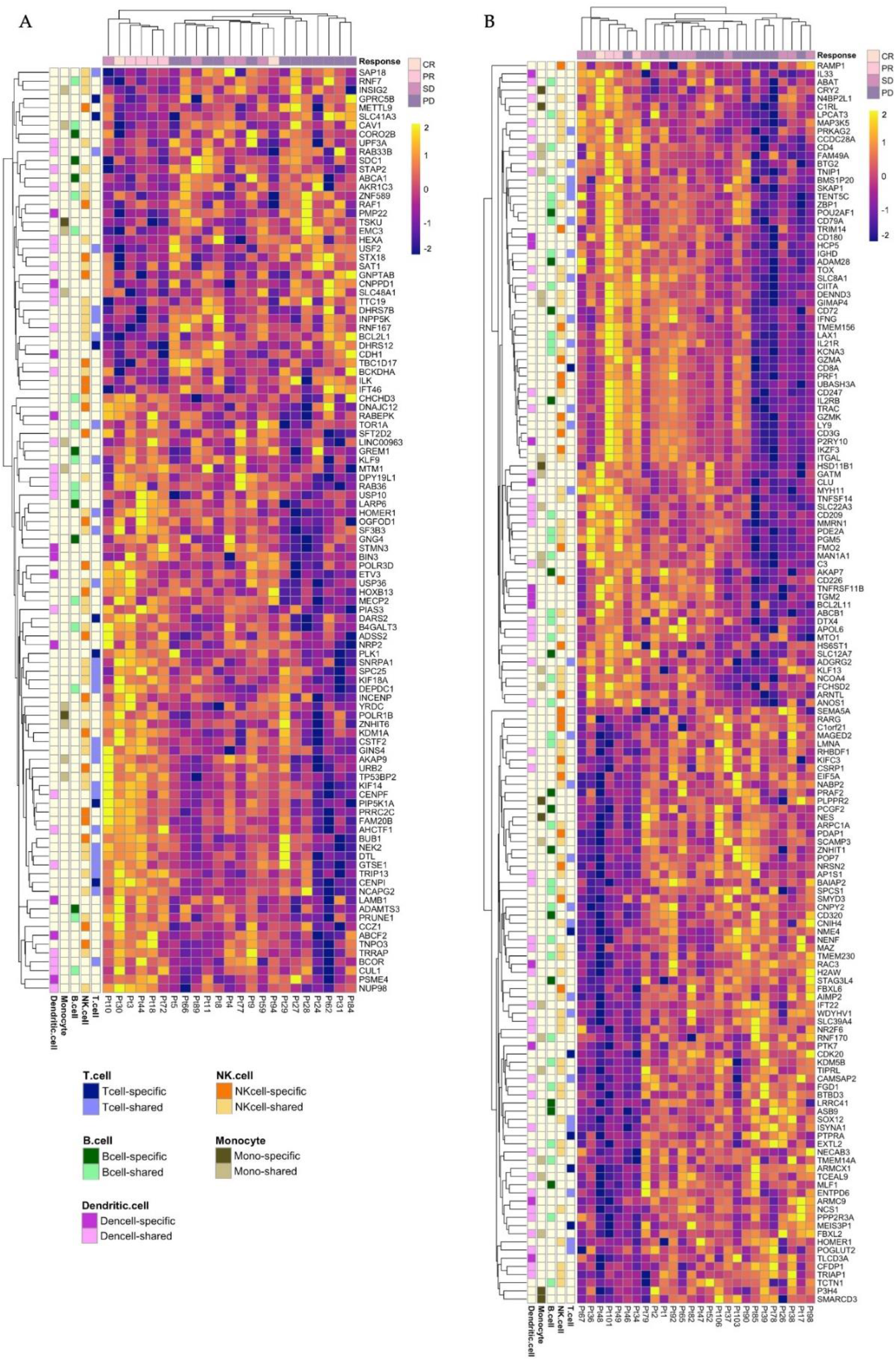
IMMUNETS genes stratify melanoma cohorts by response to nivolumab. Genes shown were found in at most two IMMUNETS networks and differentially expressed between the response groups in (A) MEL_NAI (n=23) and (B) MEL_PROG (n=26). Clinical response is shown (top) and a cluster of responsive patients is found on the left of each heatmap. The membership of genes in the IMMUNETS networks is shown at the left side of the heatmap, please see key for details. Blom-transformed gene expression is visualised on a yellow (highest) to blue (lowest) scale, thus lighter colours represent higher expression values.

### IMMUNETS: Modelling immune cell differentiation and activation

We built IMMUNETS with data from the IRIS study, which assayed key immune cell types in multiple states of activation and differentiation [21]. Each IMMUNETS network connects genes that are both differentially regulated in immune cell function and are highly correlated within the individual cell type for which the network was constructed. Differentially expressed genes were identified either between or within immune cell types and were used to construct correlation networks for T cells (T-net: 3069 genes, 54108 edges), Natural Killer (NK) cells (NK-net: 5944 genes, 65426 edges), B cells (B-net: 4626 genes, 105349 edges), Monocytes (Mono-net: 2867 genes, 55150 edges) and Dendritic cells (Dend-net: 5071 genes, 58995 edges). Together, the five correlation networks cover 9668 genes in total, of which 3262 are found in only one network and 2820 are present in exactly two of the five networks (Table 1, Supplementary Data File S1). The representation scope of IMMUNETS is defined by regulation in the 214 samples assayed by IRIS for the five cell type networks [21], and our network inference protocol. For example, the secreted cytotoxic proteases termed granzymes are key effectors of cytotoxicity found in T-net (GZMB, GZMM, GZMH) and NK-net (GZMA, GZMB, GZMK, GZMM). GZMA and GZMK are present in NK-net, which aligns with single-cell analysis of NK cells [26]; however, these genes are also important in T cell functions [27]. Indeed, genes found only in one of the five IMMUNETS networks have highly coordinated expression changes in IRIS for the corresponding cell type, but may also be expressed in other cell types. IRIS contains a further 14 samples for plasma cells, however the protocol applied here did not produce a network for these samples and therefore plasma cells are not represented in IMMUNETS. B-net has the most unique genes (n=1022) and the highest average degree, with a value of 22.8.

**Table 1.** Unique pairwise overlap between IMMUNETS cell types. Counts are shown for genes that are represented in only one (diagonal) or exactly two (off-diagonal) IMMUNETS networks. For example, the top left value (for T cell, T cell) identifies 233 genes present only in the T-net network; the value of 357, immediately below, corresponds to genes found in T-net and NK-net but not present in any other IMMUNETS network.

|  | T cell | NK cell | B cell | Monocyte | Dendritic cell |
| --- | --- | --- | --- | --- | --- |
| T cell | 233 | - | - | - | - |
| NK cell | 357 | 935 | - | - | - |
| B cell | 131 | 535 | 1022 | - | - |
| Monocyte | 58 | 171 | 120 | 233 | - |
| Dendritic cell | 143 | 750 | 333 | 222 | 839 |

NK-net is the largest overall (2748 genes) and has the greatest overlap with the other networks, for example sharing 751 and 535 genes with Dend-net and B-net, respectively. In order to refine the IMMUNETS correlation networks, we derived focus networks taking the genes from Table 1 for each immune cell type as input for the NetNC algorithm, and using HumanNet as the reference network [28,29] (Figure 2, Supplementary Figures S1, S2, S3, Supplementary Data Files S2, S3). As expected, the focus networks have connections between clusters that are annotated with biologically related terms. For example, the T cell focus network (Figure 2A) connects clusters for ‘cell cycle process’, ‘DNA repair’, ‘DNA replication’ and ‘RNA metabolic process’; similarly, there are connections in the B cell network between ‘ion transport’ and ‘transmembrane transport’ clusters.

The T cell focus network covers biological processes required for growth and proliferation (cell cycle, RNA metabolism, translation), including key T cell proliferation genes (Figure 2A). Indeed, proliferation is central to normal T cell biology [30]. A ‘regulation of lymphocyte proliferation’ (RegLP) cluster has seven genes that were found only in the T-net correlation network. The RegLP cluster includes molecules associated with p38 MAP kinase signal transduction (MAPK11, MAP2K6) [31] and cell surface glycoproteins CD8, CD28 that have well-known roles in antigen-induced activation [32]. Other cell surface glycoproteins and interleukins identified in the RegLP cluster are important for a range of T cell functions, for example: CD7 is expressed in CD8+ effector T cells [33]; CD5 downregulation potentiates T-cell antitumour activity [34]; IL4, IL13, IL5 locate in a cytokine gene cluster on chromosome five and regulate multiple immune cell functions, including in T helper 2 cells [35,36]; IL17A is produced by activated T cells where it impacts many processes, notably promoting tumour progression [37,38]. Signalling through CD3 is important for T cell activation [39]; CD3 subunits are found in multiple immune cell types in IMMUNETS but do not appear in the T cell focus network, which only contains genes from T-net and at most one other IMMUNETS network. In particular, CD3E and CD3EAP are represented in three IMMUNETS networks and therefore were not input to NetNC for focus network generation. Most of the genes in the T cell focus network (76%, 141/185) are coordinately regulated within other immune cell types in IMMUNETS, especially NK cells. On the other hand, approximately half of the genes in the B cell focus network (Figure 2B) are represented only in the B-net correlation network (51%, 409/797). These cover a wide range of cell activities, including ABC transporters, ADAM proteases, cytochrome P450 family, growth factor receptors, solute transport and many others (Supplementary Data File S2). A ‘B cell activation’ cluster contains genes involved in B cell antigen receptor complex (BCR) signaling. For example, this cluster includes CD79A, CD79B, CD19, CD22, CD20 (MS4A1), and transcription factors important for B cell development including BCL11A, PAX5 [40–45]. An ‘inflammatory response’ cluster contains IL-20 subfamily interleukins IL22 and IL22RA1, IL20RA, IL10RB which form receptor complexes that bind IL22 or IL26 [46]. IL22 is important for B cell recruitment to tertiary lymphoid structures [47] and depletion of B cells reduces IL22 production, which can stimulate cell behaviour typical of aggressive tumours [48,49]. Therefore, production of IL22 by B cells might be part of a positive feedback loop driving poor prognosis in immunoedited tumours.

### IMMUNETS genes stratify melanoma patients by response to nivolumab

We analysed the expression of immune-regulated genes found in at most two of the five IMMUNETS cell types for advanced melanoma patients [22] (Figure 3). Data were analysed from tumour biopsies taken before treatment with nivolumab, an inhibitor of PD-1 [22, 23]. Two cohorts were considered: patients previously treated with ipilimumab that had progressed (MEL_PROG, n=26) and those that were ipilimumab-naive (MEL_NAI, n=23). Both cohorts were part of the NCT01621490 trial, which took patients that were refractory, intolerant to, or had refused standard therapy [50]. MEL_NAI and MEL_PROG respectively had a total of 105 and 152 differentially expressed IMMUNETS genes between nivolumab responders and non-responders (q<0.05, 2-fold change). Therefore, we identify differentially regulated immune cell genes that correlate with response to nivolumab *in vivo*, and stratify treatment response in unsupervised clustering (Figure 3). A ‘good response’ cluster emerged in MEL_NAI and in MEL_PROG; respectively containing 0/6, 1/7 patients with progressive disease (PD). Similarly, a ‘poor response’ cluster in each cohort respectively has 1/17, 0/19 patients with either complete or partial response. Many genes have striking differences in expression between the good and poor response groups. We reasoned that immunotherapy response mechanisms would likely be represented within IMMUNETS genes that are upregulated in patients who responded to nivolumab. In contrast, a range of biological processes which drive or correlate with tumour aggressiveness could be upregulated in non-responders, and so appear a less attractive pool for discovery of candidate immunotherapy response biomarkers. Therefore, we focus on genes upregulated in tumours that respond well to therapy, because we expect that these are more pertinent to molecular control of therapy response.

Immune tolerance is mediated by expression of PD-L1 in many cell types and PD-1 in T cells, which are therefore critical in preventing autoimmune disease [51]. While nivolumab binds to and inactivates PD-1 [52], PD-L1 expression is a more informative predictor of clinical response than PD-1 [53,54]. Interestingly, results from a recent meta-analysis suggest that the PD-L1 status of tumour-infiltrating immune cells has advantages over PD-L1 staining in tumour epithelial cells for predicting ICI response [54]. PD-L1 negative patients benefited from ICI therapy in studies that examined staining in tumour cells only, whereas patients with PD-L1 negative immune infiltrate had Hazard Ratio 95% CI values overlapping unity (1.0) and so were not shown to benefit from ICI therapy. PD-1 and its ligand PD-L1 did not pass the criteria for inclusion in IMMUNETS and therefore do not appear in Figure 3. The genes found only in the T-net network and upregulated in the MEL_NAI nivolumab response group are PLK1, CENPI, PIP5K1A and DARS2. Upregulation of genes that function in cell proliferation (PLK1, CENPI) typically confers poor prognosis [55–58]. Therefore, elevated levels of PLK1 and CENPI in the nivolumab responders likely reflects expression in tumour-associated immune cells; indeed, proliferation is integral to an effective T cell immune response [59]. Correlation of immune proliferation with immunotherapy response evidences pre-existing immune activation that is enhanced by immunotherapy; aligning with reports that responsive patients have inflamed or ‘immune hot’ tumours [60]. PIP5K1A functions in phosphatidylinositol signalling which impacts multiple cell processes, including T cell activation [61], and DARS2 is a mitochondrial aspartyl-tRNA synthetase that has been suggested as a candidate causal gene in rheumatoid arthritis [62]. Thus, elevation of PIP5K1A and DARS2 in nivolumab responders is also consistent with ICI treatment acting to enhance pre-existing T cell inflammation. Genes found only in B-net that are upregulated in the MEL_NAI treatment response group were ADAMTS3, GREM1, LARP6 and GNG4. ADAMTS3 is a metalloprotease associated with Hennekam lymphangiectasia–lymphedema syndrome 3 [63]; proteases, including ADAMTS genes, are regulated in immunity and control cell-cell interactions [64]. GREM1 is a cytokine, BMP antagonist, VEGF agonist, and is downregulated in human B cells exposed to the toxin TCDD, which causes immunosuppression in addition to other harmful effects [65,66]. LARP6 is an RNA binding protein that regulates multiple aspects of cell behavior, including collagen production [67]; our analysis predicts a role in B cell function and immunotherapy response for LARP6. The presence of G protein gamma subunit GNG4 in B-net and upregulation with response to therapy suggests a role in B cell activation, consistent with the importance of G-protein coupled receptor signalling in the humoural immune response [68]. Therefore, our analysis highlights genes regulated in B cell biology and response to nivolumab in treatment-naive melanoma; the biological mechanisms involved warrant further investigation. NRP2 is one of several genes found only in Dend-net that are upregulated in the nivolumab response group. While NRP2 is expressed in multiple immune cell types, it is important for dendritic cell maturation, migration and T cell activation [69]. Overall, our results in MEL_NAI align with reports that ‘immune hot’ status correlates with response to ICIs [60] and suggest candidate nivolumab response biomarkers.

Favourable treatment response in MEL_PROG correlates with upregulation of CD247, the zeta chain of CD3, which forms a complex with the T cell receptor (TCR) and is central to the T cell immune response [39, 70]. CD247 is expressed in a wide range of leukocytes [71] and is regulated in dendritic, NK cells in IMMUNETS, rather than T cells. CD8α, produced from CD8A, is a canonical marker for the T cell population involved in tumour surveillance and forms a heterodimer with CD8β, or may homodimerise [72]; CD8A is the only gene exclusive to T-net that correlates with nivolumab response in MEL_PROG. All of the other IMMUNETS cell types had more upregulated network-specific genes in patients that respond to therapy (NK-net, 13 genes; Dend-net, 8 genes; B-net, 6 genes; Mono-net, 3 genes). These include: from NK-net, the CD226 cell surface glycoprotein that regulates NK cell antitumour activity [73]; from B-net, the POU2AF1 transcriptional factor essential for B cell maturation [74] and ADAM28, which controls B cell transendothelial migration [75]; from Mono-net, the hydroxysteroid 11β dehydrogenase type 1 (HSD11B1) that regulates resolution of the inflammatory response, including in macrophages, and correlates with CD4+ T cell activation [76]; from Dend-net, IL33 that stimulates CD8+ T cell antitumour responses and downregulates PD-1 [77]. Our analysis with IMMUNETS suggests that dendritic cells could be a source of IL33 in antitumour immunity; although IL33 is produced by other cell types [78]. Several genes upregulated in the MEL_PROG good response group can drive tumour aggressiveness when expressed in cancer cells. For example, ADAM28 expression in tumour epithelium associates with poor prognosis, but is important for T cell mobilisation to metastatic lesions [79,80]. Therefore, upregulation of otherwise poor prognosis genes in the good response group suggests that their correlation with ICI therapy response arises from regulation within immune cells.

DAVID [80] analysis of immune-regulated genes that are differentially expressed between therapy response groups in MEL_NAI and MEL_PROG (Table 2, Supplementary Tables S1, S2) was consistent with results from NetNC and the heatmap hierarchical clustering. Broadly, MEL_NAI has significant functional clusters for processes involved in proliferation, while the differences in treatment response for MEL_PROG associate with immune regulation, including T cell signalling and activation. As noted above, proliferation genes correlate with good response to treatment in MEL_NAI, are regulated in IMMUNETS and therefore may reflect immune cell proliferation. The difference between mechanisms of response in the two cohorts is underlined by finding only one IMMUNETS gene significantly changed in both cohorts; HOMER1, a scaffolding and signal transduction protein that regulates T cell activation [82,83]. Moreover, the expression level of HOMER1 has an opposite relationship with clinical response MEL_NAI and MEL_PROG. These differences could potentially arise from expression of different functional isoforms, [83], or HOMER1 membership of different protein signalling complexes in MEL_NAI and MEL_PROG.

**Table 2.**
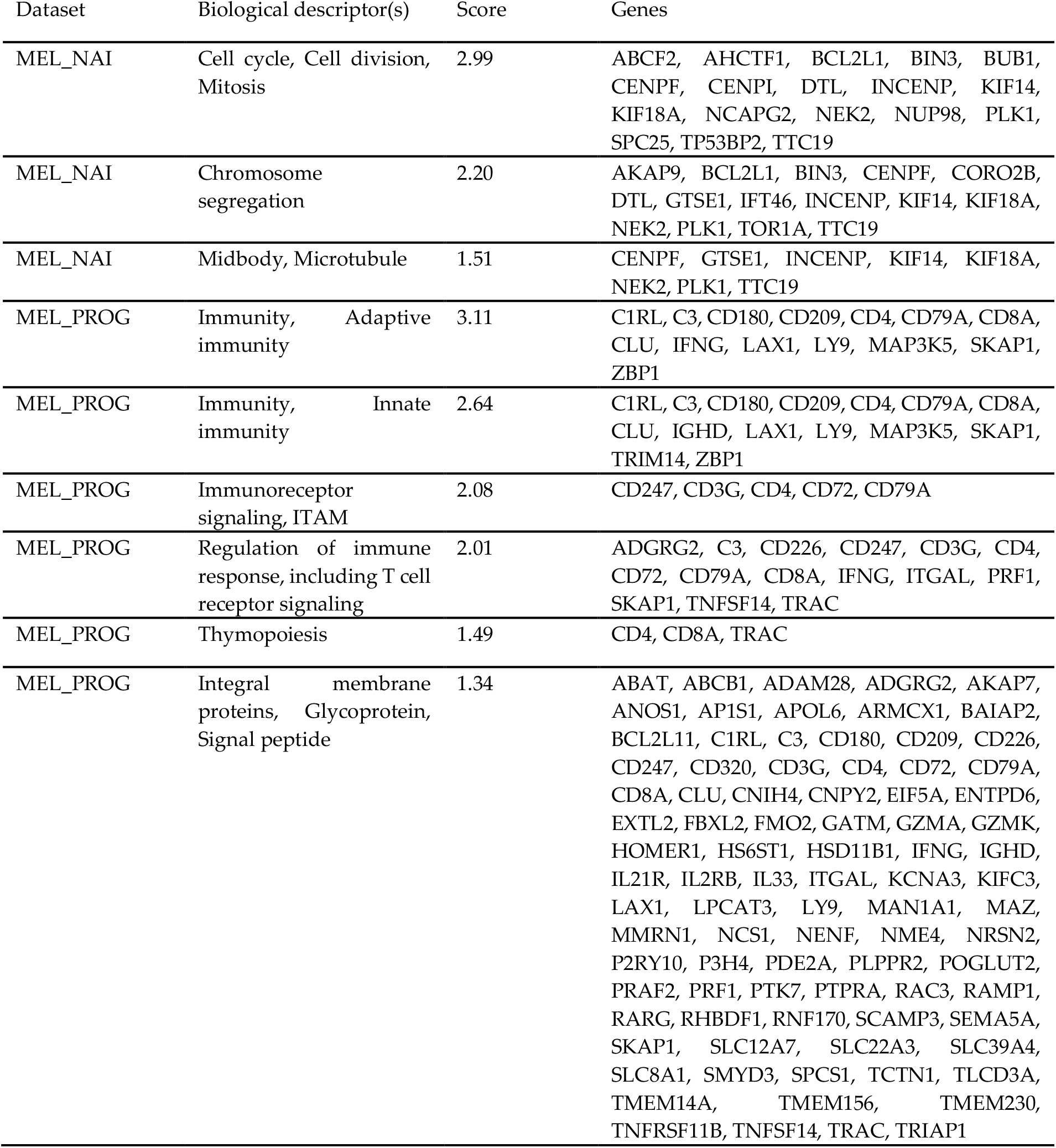
Summary of DAVID clusters for immune-regulated, differentially expressed genes in MEL_NAI (n=105) and MEL_PROG (n=152). Clusters with significant enrichment score (≥1.3) are shown.

### Candidate immune biomarkers risk stratify independent melanoma and ovarian carcinoma cohorts by overall survival

We sought to identify candidate ICI response biomarkers from genes that are coregulated in IMMUNETS because they may capture the status of tumour-associated immune cells, which is the primary substrate for ICI therapy. We took IMMUNETS genes that are upregulated in patients who responded to nivolumab because these may be superior candidate treatment response biomarkers than genes that are upregulated in non-responders. Indeed, a proportion of the genes associated with poor response to nivolumab may act within biological mechanisms that drive cancer progression in multiple contexts, rather than directly influencing the response to therapy. Many of the IMMUNETS genes that are differentially expressed in MEL_NAI function in proliferation and the cell cycle; values for these genes may measure expression for tumour cells in the biopsied material. Activation of proliferation in tumour epithelium is typically a poor prognosis factor [84] and so inclusion of cell cycle or proliferation genes does not align well with our aim to study immune regulation in ICI therapy response. Accordingly, we prioritised candidate biomarkers for further study according to either: a) membership of a significant immune-related functional annotation cluster (Table 2); b) presence in only one of B-net, T-net or Mono-net and upregulation in the MEL_PROG nivolumab response group (Figure 3); c) from prior knowledge of ICI response factors (PD-L1, PD-1 TGFB1, TGFBR2) [53,54,85]; comprising a total of 27 genes (BIO_27).

We investigated the prognostic value of BIO_27 in TCGA data from melanoma (n=390, MEL_TCGA) and ovarian carcinoma (n=375, OV_TCGA) [24,25] with univariate analysis of patient groups defined using unsupervised clustering of BIO_27 expression values. Following correction for multiple hypothesis testing [86], 22/27 MEL_TCGA genes and 2/27 OV_TCGA genes are significant (log-rank *q*<0.05). Notably, increased expression of ZBP1 and CD79A confers good prognosis in both MEL_TGCA and OV_TCGA (Table 3, Supplementary Figures S4, S5). ZBP1 is a cytosolic DNA sensor that induces type I-interferon production in innate immunity [87,88] and enhances the T cell response in melanoma vaccine studies [89,90]. These functions align well with our results suggesting a role for ZBP1 in enhancing immunotherapy response (Figure 3) and in contributing to tumour clearance (Table 3). CD79A heterodimerises with CD79B to form the signal transducing component of the B cell antigen receptor complex [91] and demarcates B cells in single cell analysis [92]. Thus, our findings suggest that CD79A expression may quantify B cell tumour infiltration and antitumour immunity in melanoma and ovarian carcinoma. Additional B cell genes are prognostic in MEL_TGCA, including CD72, a BCR co-receptor that is important for B cell proliferation, differentiation and survival [93]. Therefore, in contrast to previous analysis based on signalling effectors LCK and SYK [24], we identify prognostic value for both T cells and B cells in MEL_TCGA. Recruitment criteria ensured that MEL_TCGA comprised only patients with no previous systemic treatment, excepting adjuvant interferon ≥90 days prior, and previous work reports a correlation between immune activation and good prognosis in this cohort [24]. In agreement with these findings, ten of the thirteen (77%) selected IMMUNETS genes that are upregulated in MEL_PROG therapy response group also portend good prognosis in our analysis of MEL_TCGA (Table 3). Therefore, we delineate factors that are regulated in immune cells, which correlate with response to ICI therapy and may drive tumour clearance in other melanoma treatment pathways.

**Table 3.**
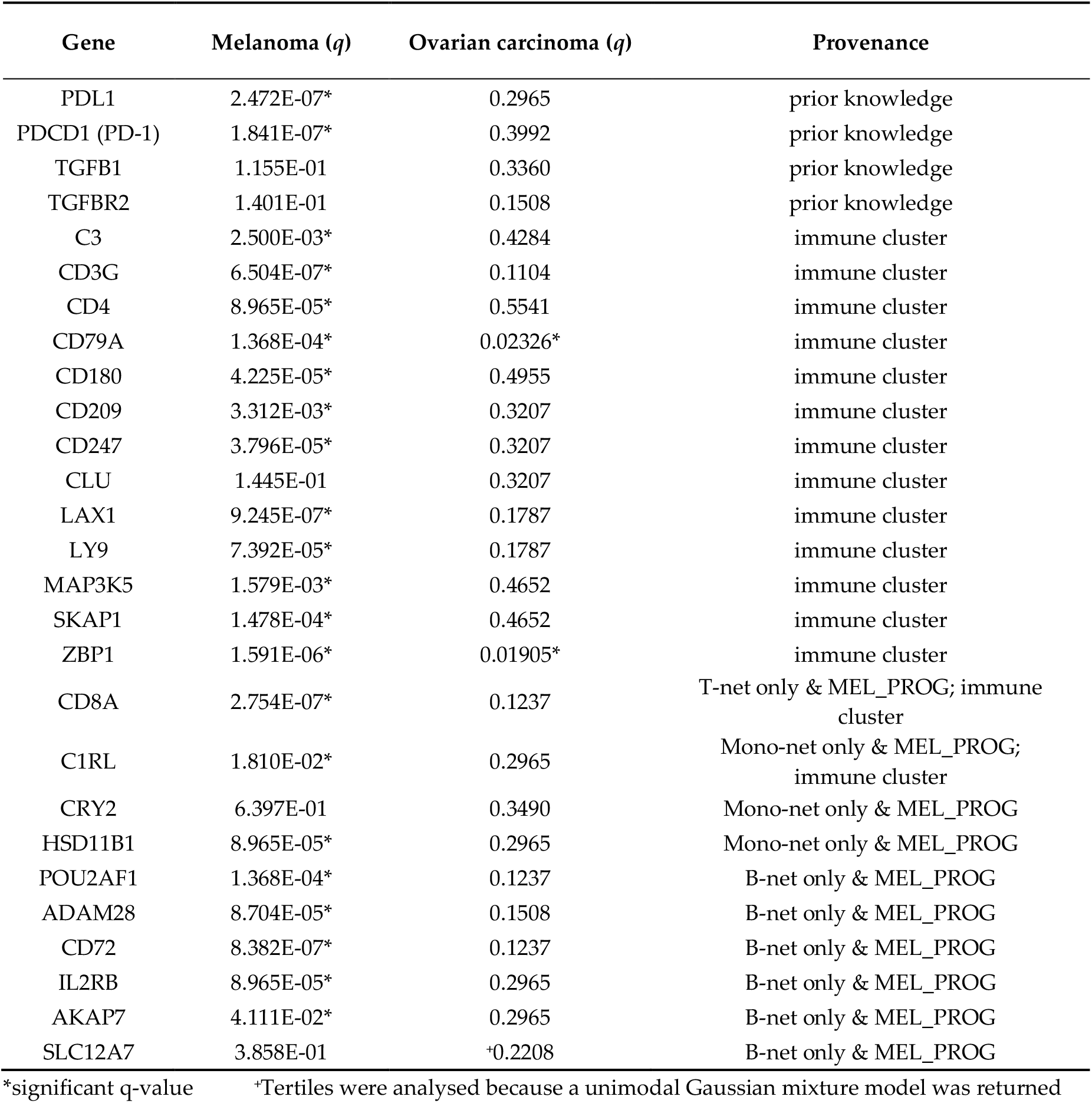
Univariate risk stratification of melanoma and ovarian carcinoma by overall survival with BIO_27. Log-rank test q-values are shown, with adjustment for multiple hypothesis testing. The provenance column indicates genes selected for univariate analysis according to prior knowledge; from an immune functional annotation cluster (Table 2); genes only in T-net, B-net or Mono-net and upregulated in MEL_PROG (Figure 3).

Stepwise feature selection with Bayesian information criterion (BIC) regularisation was performed for Cox proportional hazards modelling of overall survival, taking as input the genes that were significant in univariate analysis, as well as age and tumour stage [94,95]. Significant models were returned for MEL_TCGA (tumour stage, age, CD72; LR *p*<10^-9^) and MEL_OV (age, CD79A; LR *p*<2×10^-5^). All of the selected features were individually significant in the multivariate models (Table 4, Table 5). Regularisation with BIC ensures that the selected molecular features add information over and above the clinical variables analysed. As expected, higher tumour stage and age confer worse prognosis. The melanoma model satisfies the proportional hazards assumption, whereas the ovarian model does not (Supplementary Table S3). Both CD72, CD79A correlate with lower risk (respective HR 0.90, 0.97), are B cell markers [91–93] and were found only in B-net. Therefore, these results highlight the importance of B cells and humoural immunity in controlling melanoma and ovarian carcinoma progression. Indeed, the differences in B cell markers between nivolumab response groups, including BCR signalling, (Figure 3) align with growing evidence that this cell type plays an important role in successful ICI therapy [96].

**Table 4.**
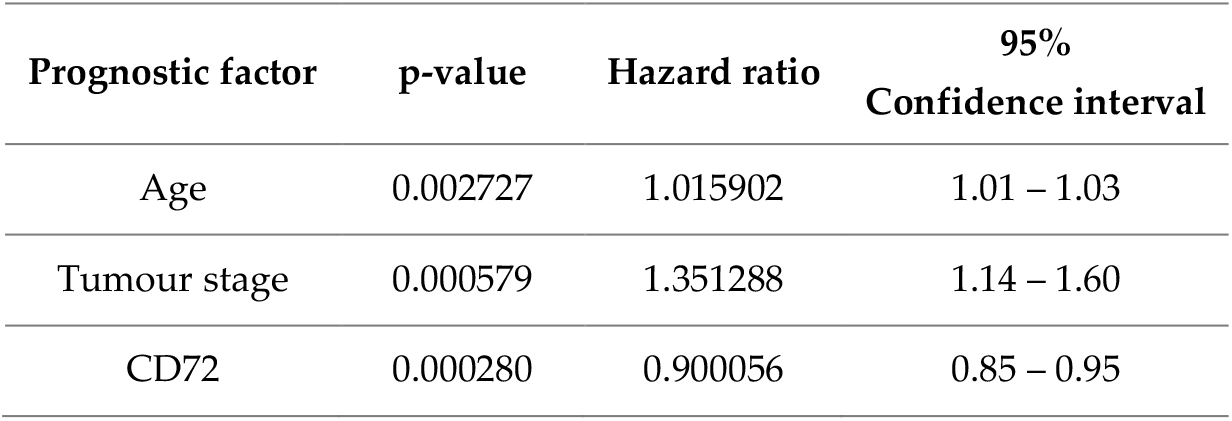
Regularised Cox proportional hazards model for melanoma.

**Table 5.**
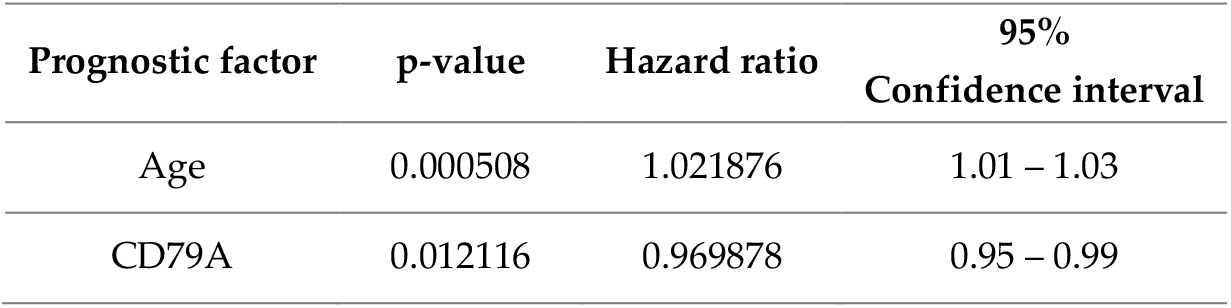
Regularised Cox proportional hazards model for ovarian carcinoma.

| Prognostic factor | p-value | Hazard ratio | 95%<br>Confidence interval |
| --- | --- | --- | --- |
| Age | 0.000508 | 1.021876 | 1.01 – 1.03 |
| CD79A | 0.012116 | 0.969878 | 0.95 – 0.99 |

## Discussion

We present five immune cell networks (IMMUNETS) which model gene coregulation in T cells (T-net), B cells (B-net), NK cells (NK-net), monocytes (Mono-net) and dendritic cells (Dend-net) (Table 1, Supplementary Data File 1). IMMUNETS provide a resource for understanding the pathways and protein complexes that control immune cell activation and differentiation, including systems immunology approaches [97]. Our approach is complementary to tissue-based resources such as ImSig [98]. We took IMMUNETS genes that are coregulated in at most two cell types as a basis for derivation of immune cell focus networks, which capture key biological processes regulated within each immune cell type, including immune-specific gene clusters (Figure 2, Supplementary Data Files S2, S3). We find that regulation of proliferation is a major theme within the T cell focus network, where 76% of genes are represented within other immune cell networks. In contrast, the majority of genes in the B cell focus network are not covered by another IMMUNETS cell type; a wide range of cell functions found only in B-net are present in the B cell focus network, including BCR complex signaling.

Analysis with IMMUNETS genes revealed correlates of nivolumab response in two advanced melanoma cohorts (Figure 3). The MEL_PROG cohort had progressed on ipilimumab, whereas MEL_NAI had not received prior immune checkpoint therapy [22]. The biological mechanisms underlying nivolumab response differ between the two cohorts, clearly demonstrated in functional clustering (Table 2); MEL_NAI immunotherapy response genes broadly function in proliferation, while the response in MEL_PROG is characterized by immune regulation. Genes that associate with poor prognosis when expressed in epithelial tumour cells, for example drivers of proliferation, were upregulated in patients who responded well to nivolumab; therefore, these genes are likely to mark immune cell proliferation in our analysis. Our results illuminate different tumour microenvironments between ipilimumab-naive and ipilimumab progressing patients, as well as in differing ICI response trajectories. We focus on genes upregulated in patients who responded to nivolumab, in order to help obtain a clearer readout for the cellular machinery driving response to therapy. CD8A, crucial for tumour immunosurveillance [32], is the only gene specific to T-net and elevated in the MEL_PROG response group; the corresponding set of genes from MEL_NAI function in proliferation and T cell activation, consistent with reports that T cell inflamed or ‘immune hot’ tumours are more responsive to ICIs [60]. Overall, our analysis suggests greater importance of T cell biology in regulating MEL_NAI response to therapy, compared with MEL_PROG. Relative to T cells, we find larger numbers of nivolumab response genes that are specific to NK-net or Dend-net. Indeed, many IMMUNETS genes regulated in Dendritic cells and NK cells differentiate therapy response groups in both MEL_NAI and MEL_PROG, highlighting the importance of these cell types in activation of antitumour immunity with checkpoint inhibitors. One example is IL33 which is only found in Dend-net and therefore is regulated by dendritic cell stimulation by LPS [21,99]; IL33 is important for inflammation including in dendritic cell dependent stimulation of T cell antitumour activity [77]. Therefore, this analysis suggests that IL33 might drive a positive feedback loop involving activation of dendritic cells, T cell immunity and ICI response. Multiple B cell genes are also differentially expressed between nivolumab response groups. These results offer a window into the complex interplay between multiple immune cell types in ICI therapy response and resistance.

IMMUNETS genes that correlate with MEL_PROG therapy response risk stratify melanoma [24] and ovarian carcinoma [25] cohorts by overall survival (Tables 3-5). Therefore, our analysis of independent cohorts pinpoints immune-regulated genes that correlate with both ICI response and overall prognosis; these genes may be candidate biomarkers for ICI therapy response and warrant further investigation. All five immune cell types studied are represented in the significant genes from univariate analysis, in line with the importance of dynamic cell-cell communication and higher order cooperativity in antitumour immunity. Multivariate analysis with regularised feature selection highlights the prognostic value of CD72 and CD79A, controlling for tumour stage and age. Interestingly, both of these genes are B cell markers [91–93], were found only in B-net, correlate with good prognosis and are upregulated in the MEL_PROG nivolumab response group; adding to evidence that multiple cell immune types regulate checkpoint inhibitor response [96].

## Methods

### Co-expression gene networks and focus network construction

The Immune Response In Silico (IRIS) immune cell gene expression data was obtained from the Gene Expression Omnibus database, accession GSE22886 [21]. ANOVA with Benjamini-Hochberg false discovery rate correction was applied to determine significantly differentially expressed genes across the full IRIS dataset (*q*<0.01) [86,100]. Pearson correlation was calculated separately for each leukocyte dataset and only highly correlated significant gene pairs were retained (r>0.9, *q*<0.01). Therefore, the correlation values reflect co-regulation within, rather than between, each of the five cell types. Correlation-based distance measures perform well in separating functionally related genes from randomly selected pairs [101]. Genes from each of the five correlation networks for the regulated genes were taken as input to the NetNC algorithm if they were found in no more than two IMMUNETS networks; in order to produce five focus networks. NetNC analysis used the FTI setting and HumanNet as the base network [28,29]. The five focus networks output from NetNC-FTI were visualized by Cytoscape and annotated with the BiNGO plugin using a significance threshold of *q*<0.05 [102,103] (Supplementary Data File S3), all of the expressed genes in IRIS [21] were taken as the background gene list for enrichment analysis.

### Differential expression analysis of IMMUNETS genes in melanoma treatment response

RNA-seq data from melanoma patients who received treatment with nivolumab were obtained as FPKM values from the Gene Expression Omnibus database, accession GSE91061 [22]. We analysed forty-nine patients with complete data, in two cohorts: individuals without prior immune checkpoint inihbitor treatment (MEL_NAI, n=23) and those who had received Ipilimumab but had progressed (MEL_PROG, n=26). The response of patients to nivolumab was defined by the Response Evaluation Criteria in Solid Tumors (RECIST) version 1.1. Complete response (CR) and partial response (PR) indicate the elimination or decrease of lesion; stable disease (SD) indicates no significant increase or decrease; and progressive disease (PD) represents significant tumour growth [104]. We classified CR and PR as ‘response’ and PD as ‘non-response’; patients with SD were not assigned to either group. The 5898 IMMUNETS genes that were shared by no more than two of the five cell types and measured in the melanoma RNA-seq data [22] were taken forwards into differential expression analysis between the response and non-response groups. Differential expression was assessed separately in MEL_NAI and MEL_PROG (Mann-Whitney *q*<0.05, 2-fold change), with appropriate false discovery rate correction [86]. Expression values for the differentially expressed genes were Blom transformed before unsupervised hierarchical clustering with Euclidean distance and visualisation as a heatmap [105]. Differentially expressed genes for MEL_NAI and MEL_PROG were analysed separately using DAVID with the 5898 IMMUNETS genes (above) as the reference gene list, Functional Annotation Clusters with enrichment score >1.3 were taken as significant [81].

### Evaluation of candidate biomarkers for risk stratification of melanoma and ovarian carcinoma

For univariate analysis, risk groups were defined by Gaussian mixture modelling (GMM) with unsupervised selection of cardinality [106] using the BIO_27 gene expression values in MEL_TGGA or OV_TGGA; except for SKAP1 in OV_TCGA where GMM returned a unimodal model and so tertiles were taken. Log-rank test p-values with false discovery rate correction [86] identified significant genes (*q*<0.05). Multivariate analysis was conducted with genes that were significant in univariate analysis along with tumour stage and age. Tumour stage was coded numerically, translating from the ordinal values of the TNM staging system for melanoma and the FIGO staging system for ovarian carcinoma [107,108]. Stages I, Ia, Ib, Ic were assigned a value of 1; stages II, IIa, IIb, IIc a value of 2; stages III, IIIa, IIIb, IIIc a value of 3; and stage IV was assigned a value of 4. We selected features for Cox proportional hazards modelling using stepwise backwards elimination and Bayesian Information Criterion (BIC) regularisation [94,95]. The proportional hazards assumption was evaluated using the Grambsch-Therneau test [109] (Supplementary Table S3).

## Acknowledgments

Thanks to Chang Kim, Guillaume Stewart-Jones and Helen Coleman for helpful discussion.

## Author Contributions

Conceptualization, I.M.O.; Data curation, D.V., G.M., I.M.O.; Formal analysis, D.V., G.M., I.M.O.; Funding acquisition, I.M.O.; Investigation, D.V., G.M., I.M.O., Methodology, D.V., I.M.O. Project administration, I.M.O.; Resources, I.M.O.; Supervision, I.M.O.; Visualization, D.V., I.M.O. Writing—original draft preparation, D.V., I.M.O. Writing—review and editing, D.V., G.M., I.M.O. All authors have read and agreed to the published version of the manuscript.

## Supplementary Information

Figure S1: Natural Killer (NK) cell focus network, Figure S2: Monocyte focus network, Figure S3: Dendritic cell focus network, Figure S4: Survival analysis of BIO_27 in melanoma, Figure S5: Survival analysis of BIO_27 in ovarian cancer, Table S1: Functional Annotation Clustering result for MEL_NAI, Table S2: Functional Annotation Clustering result for MEL_PROG, Table S3: Grambsch–Therneau test of the proportional hazards assumption, Supplementary Data File S1: IMMUNETS networks, Supplementary Data File S2: IMMUNETS focus networks Cytoscape session, Supplementary Data File S3: BiNGO results for IMMUNETS focus networks.

