## Supplementary Information for "Immune cell networks uncover candidate biomarkers of cancer immunotherapy response"

**Supplementary Information for Vo D, McGleave G & Overton I. 'Immune cell-specific networks uncover candidate markers of cancer immunotherapy response'**

Page

Supplementary Figures:

Figure S1. Natural Killer (NK) cell focus network …………………………………………………… 2

Figure S2. Monocyte focus network …………….…………………………………………………… 3

Figure S3. Dendritic cell focus network ..………….……………………………………………… 4

Figure S4. Survival analysis of BIO_27 in melanoma ………………………………………………  5-9

Figure S5. Survival analysis of BIO_27 in ovarian cancer ….………………………………………  10-14

Supplementary Tables:

Supplementary Tables S1 and S2 are provided separately.

Table S3. Grambsch–Therneau test of the proportional hazards assumption ……………………….. 15

Supplementary Data (not included in this document):

Supplementary Data File S1. IMMUNET networks.

Supplementary Data File S2. IMMUNET focus networks Cytoscape session.

Supplementary Data File S3. BiNGO results for IMMUNET focus networks.

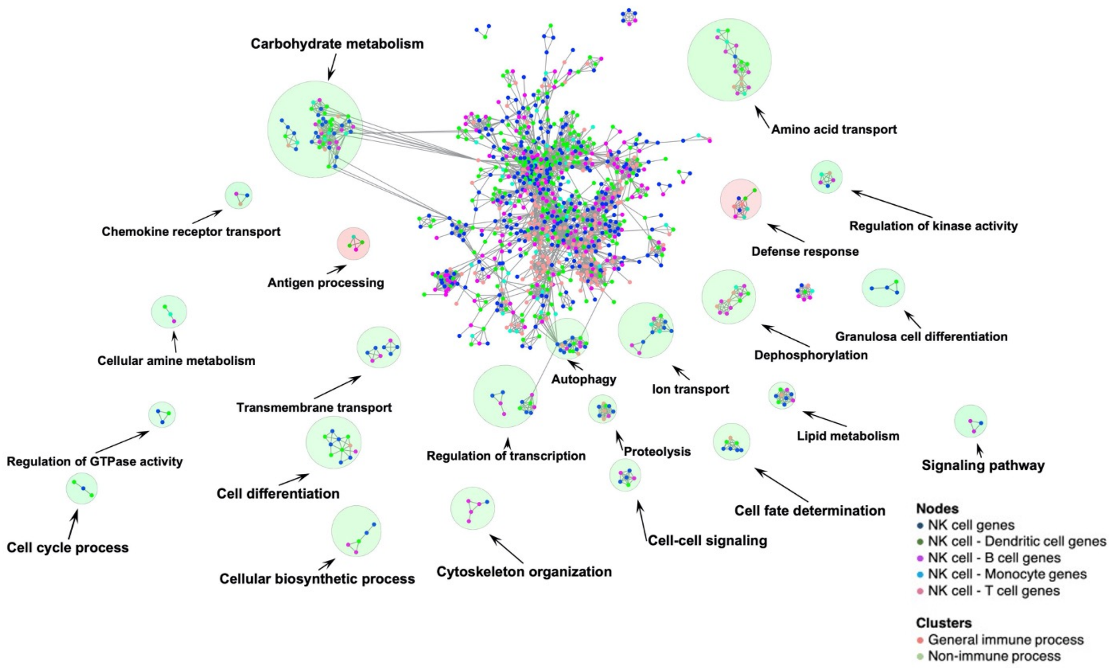


**Figure S1. Natural Killer (NK) cell focus network.** Results are shown from NetNC analysis of genes found only in NK-net, or in NK-net and one other IMMUNET network. Node colour indicates the representation of each gene in IMMUNET, according to the key. For example, genes shown in blue are only present in NK-net. Clusters are annotated according to significant terms using the BiNGO plugin. Clusters circled in red have immune process annotations, specifically "Antigen processing" and "Defense response". Clusters circled in green are annotated with GO terms that are not specific to immune cells.

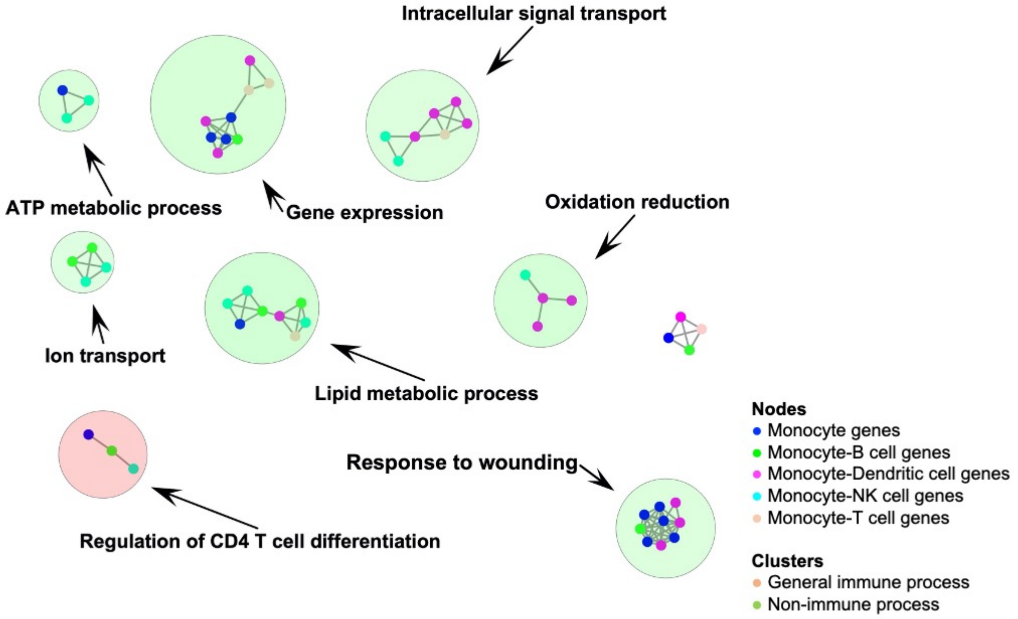


**Figure S2. Monocyte focus network.** Results are shown from NetNC analysis of genes found only in Mono-net, or in Mono-net and one other IMMUNET network. Node colour indicates the representation of each gene in IMMUNET, according to the key. For example, genes shown in blue are only present in Mono-net. Clusters are annotated according to significant terms using the BiNGO plugin. Clusters circled in red have immune process annotations, specifically "Regulation of CD4 T cell differentiation" and "Response to stress and wounding". Clusters circled in green are annotated with GO terms that are not specific to immune cells.

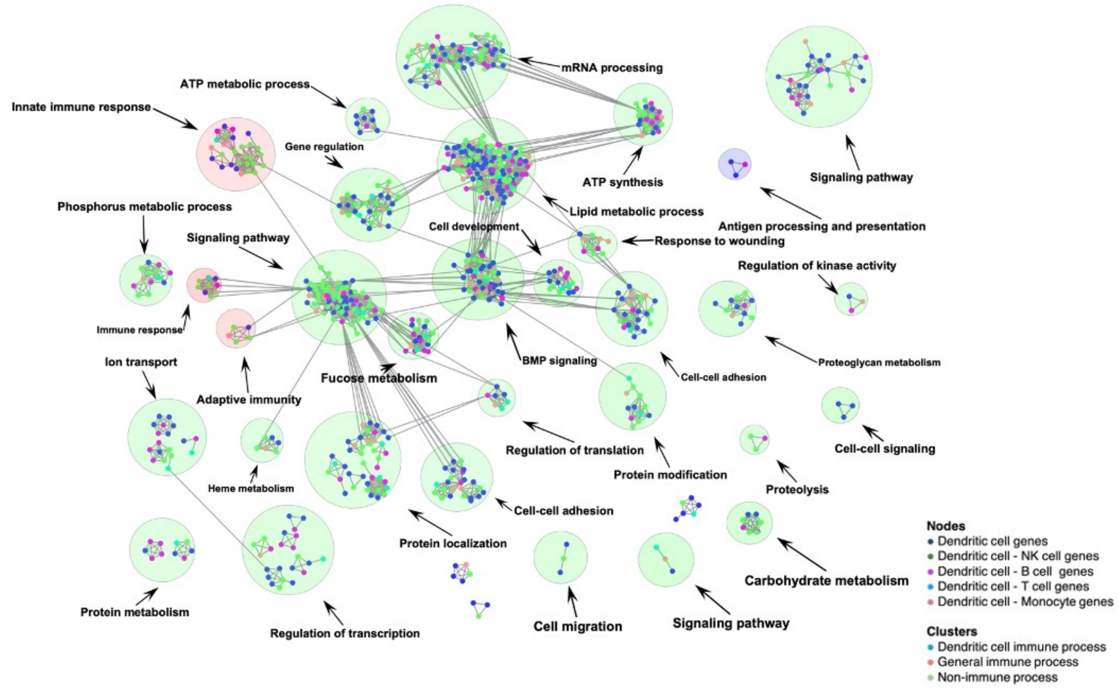


**Figure S3. Dendritic cell focus network.** Results are shown from NetNC analysis of genes found only in Dend-net, or in Dend-net and one other IMMUNET network. Node colour indicates the representation of each gene in IMMUNET, according to the key. For example, genes shown in blue are only present in Dend-net. Clusters are annotated according to significant terms using the BiNGO plugin and clusters with three more genes are shown. The clusters circled in red have immune process annotations, specifically “antigen processing and presentation”, “adaptive immunity” and “immune response” and a cluster circled in blue . Clusters circled in green are annotated with GO terms that are not specific to immune cells.

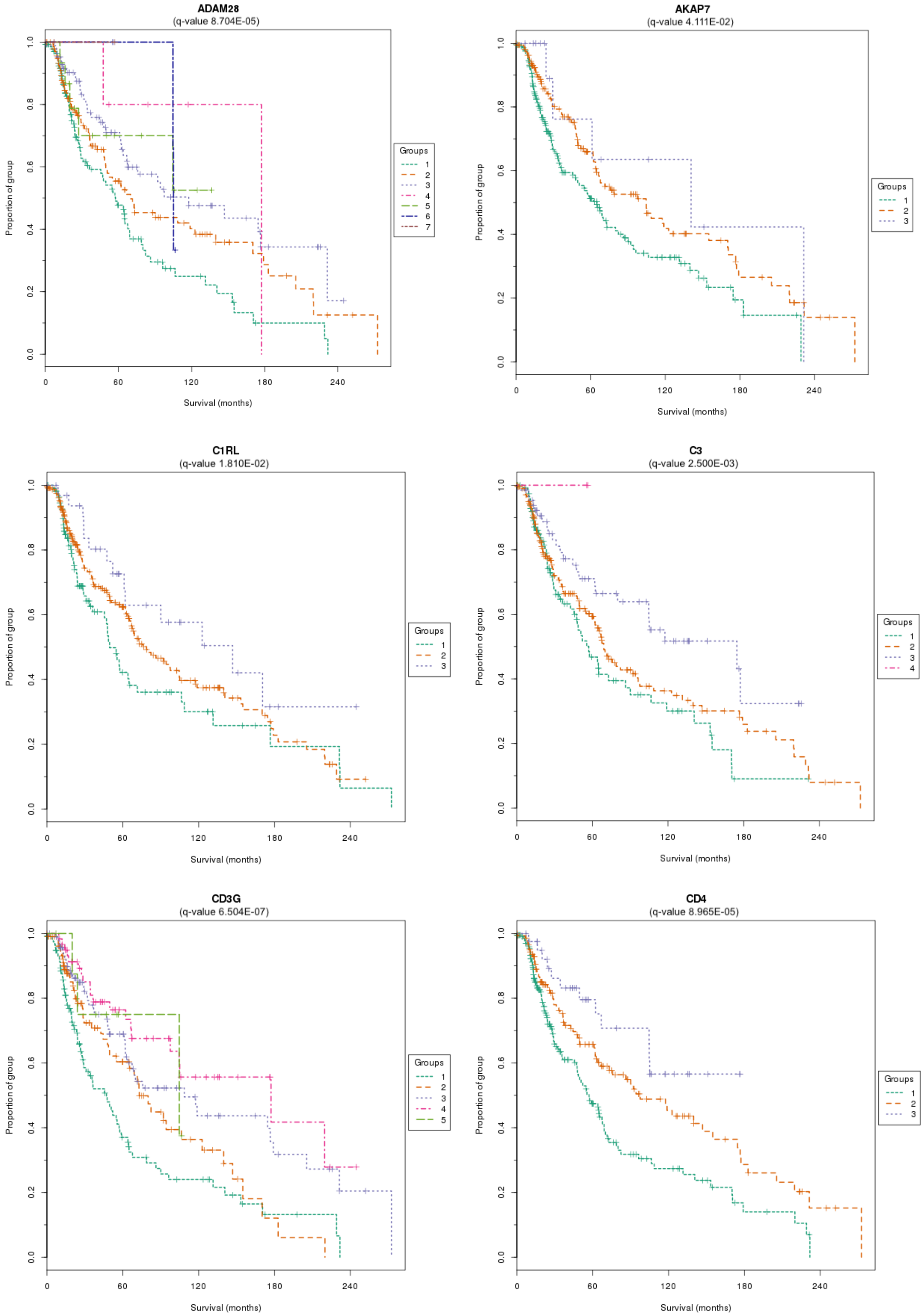


**Figure S4. Survival analysis of BIO_27 in melanoma.** Continued on the next page.

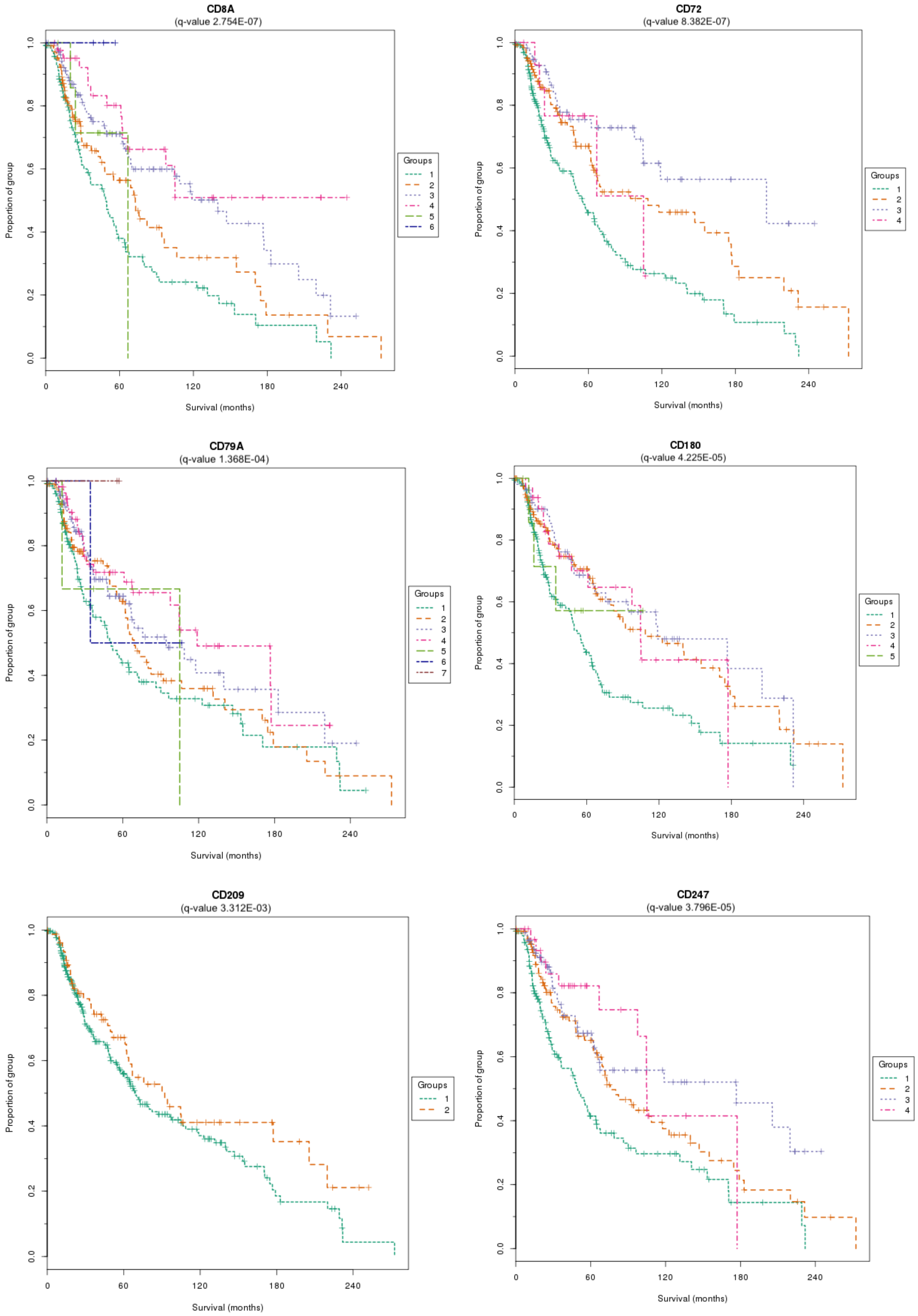


**Figure S4. Survival analysis of BIO_27 in melanoma.** Continued on the next page.

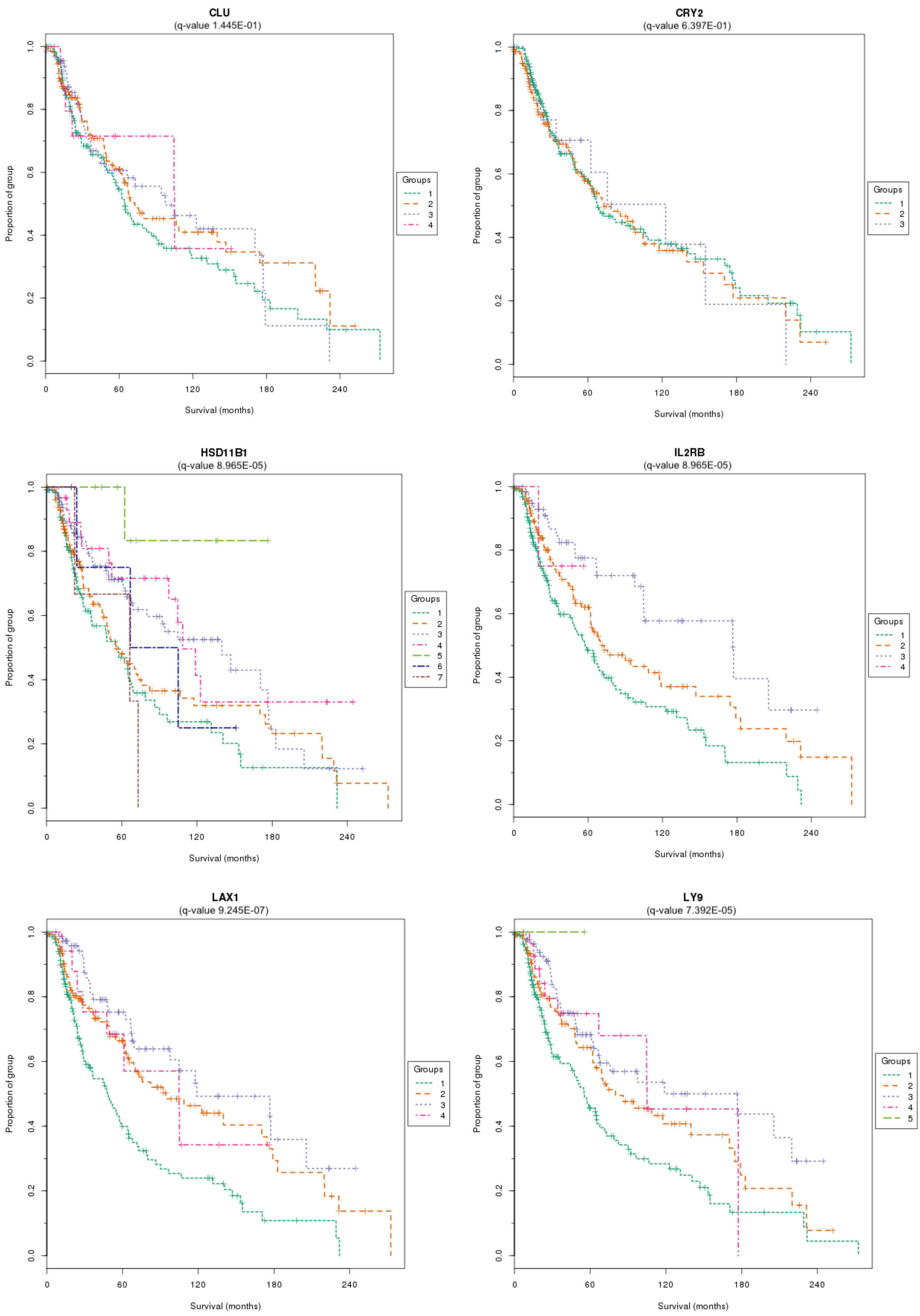


**Figure S4. Survival analysis of BIO_27 in melanoma.** Continued on the next page.

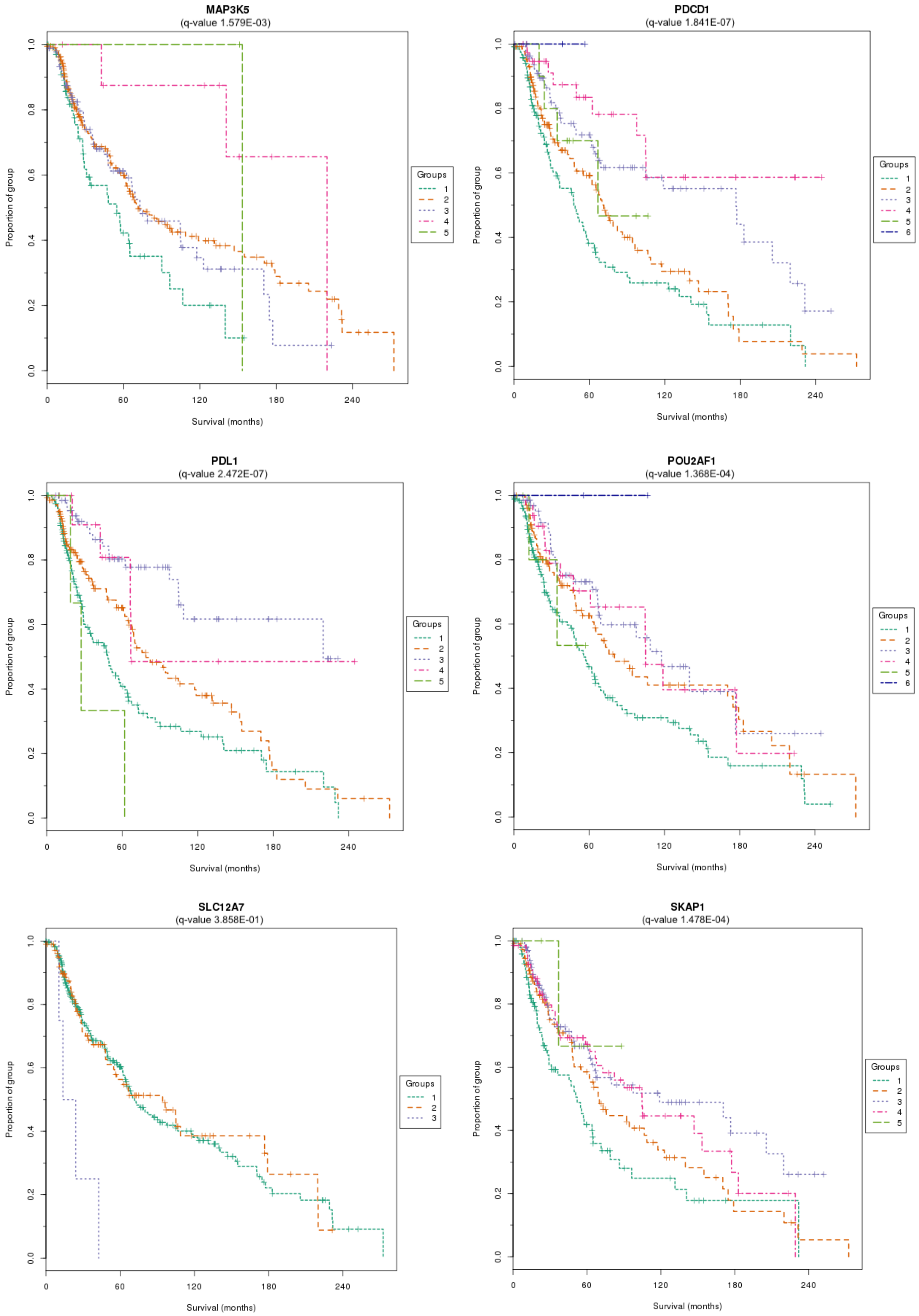


**Figure S4. Survival analysis of BIO_27 in melanoma.** Continued on the next page.

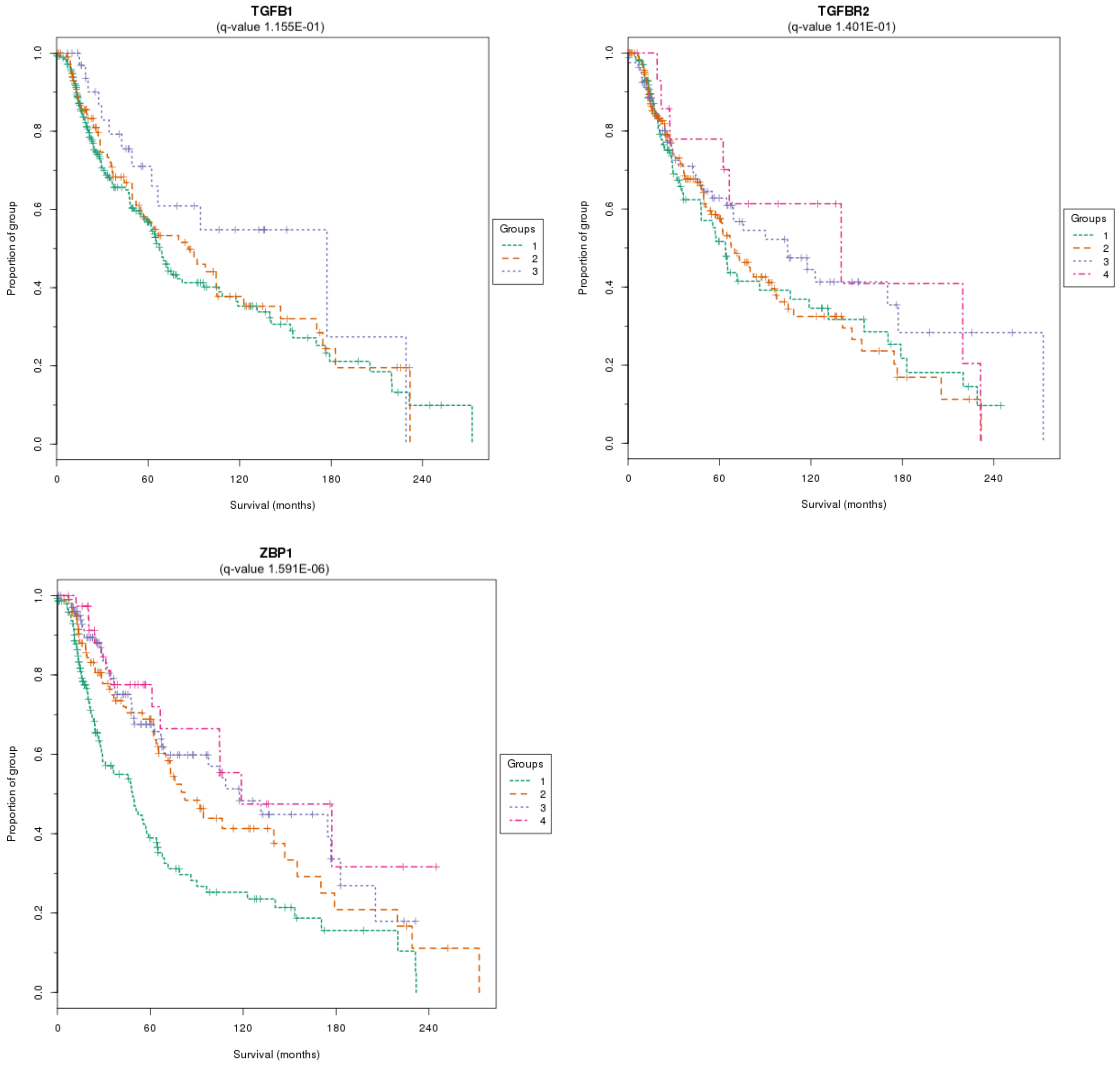


**Figure S4. Survival analysis of BIO_27 in melanoma.** Groups were defined by Gaussian Mixture Modeling of genes expression values in cutaneous melanoma patients from TCGA (n = 390). Numbering in the key identifies each sub-group; higher numbering corresponds to a higher expression level of the gene analysed. Log-rank q-values are given for each gene. Also see Table 3 in the main manuscript.

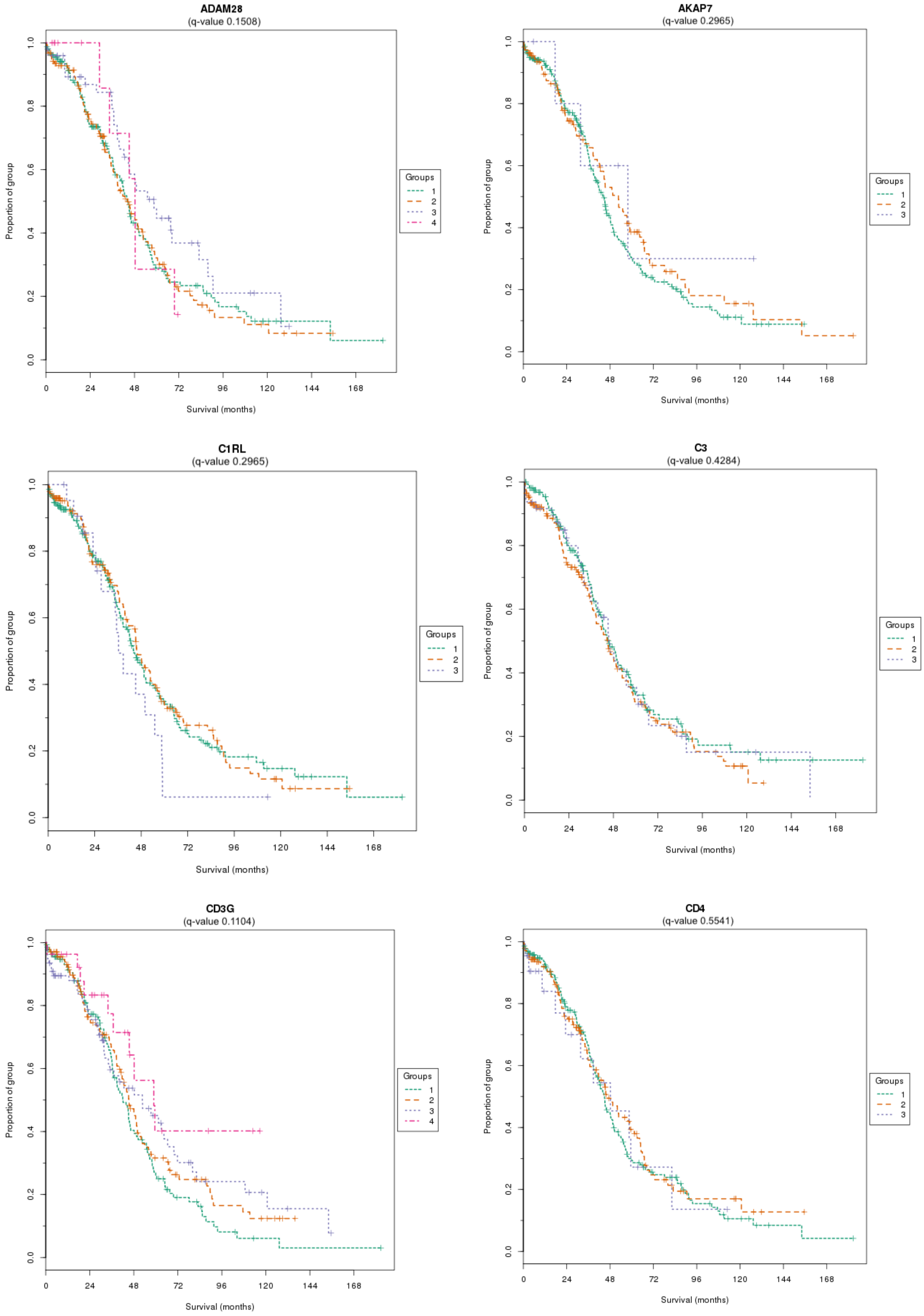


**Figure S5. Survival analysis of BIO_27 in ovarian cancer.** Continued on the next page.

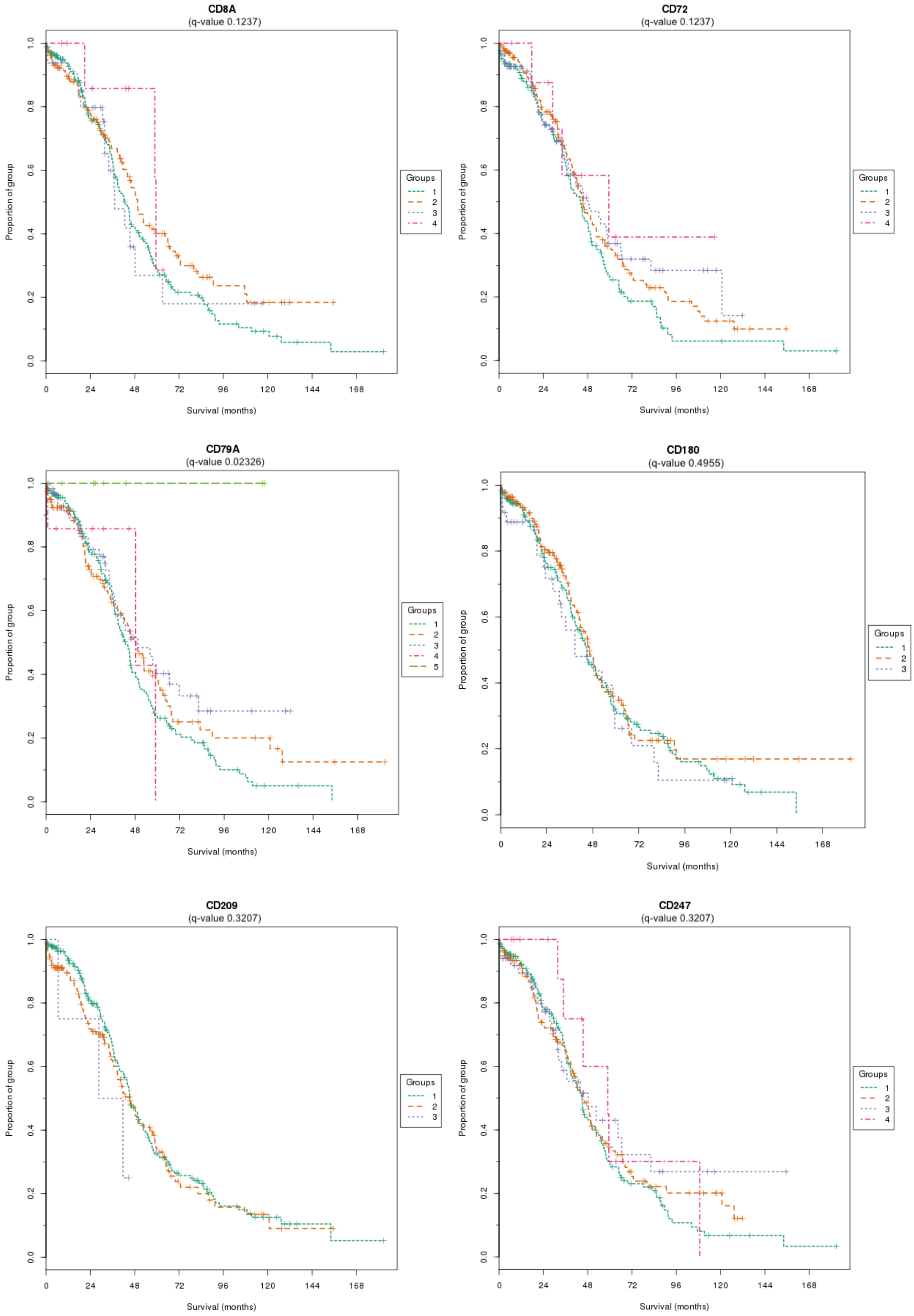


**Figure S5. Survival analysis of BIO_27 in ovarian cancer.** Continued on the next page.

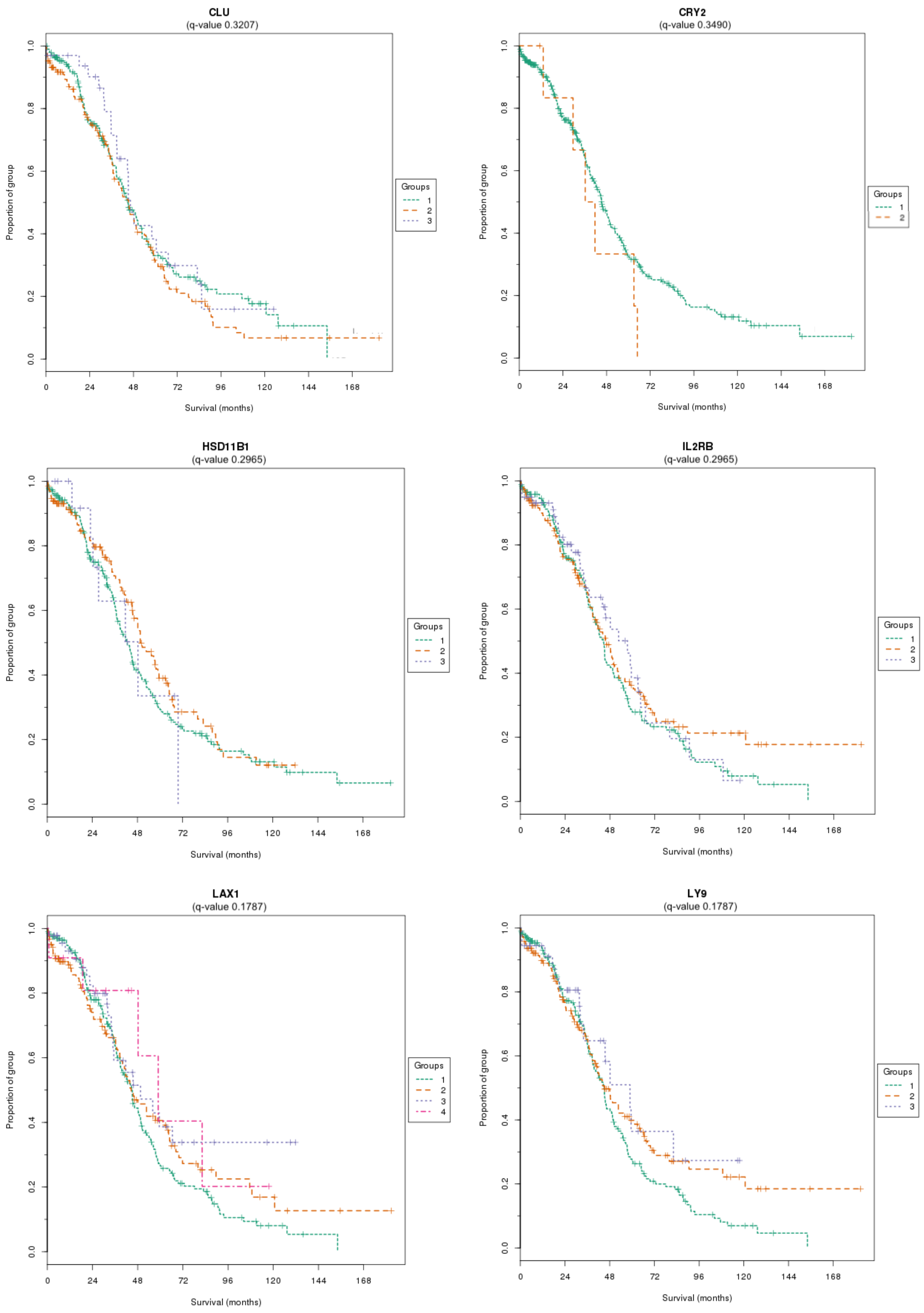


**Figure S5. Survival analysis of BIO_27 in ovarian cancer.** Continued on the next page.

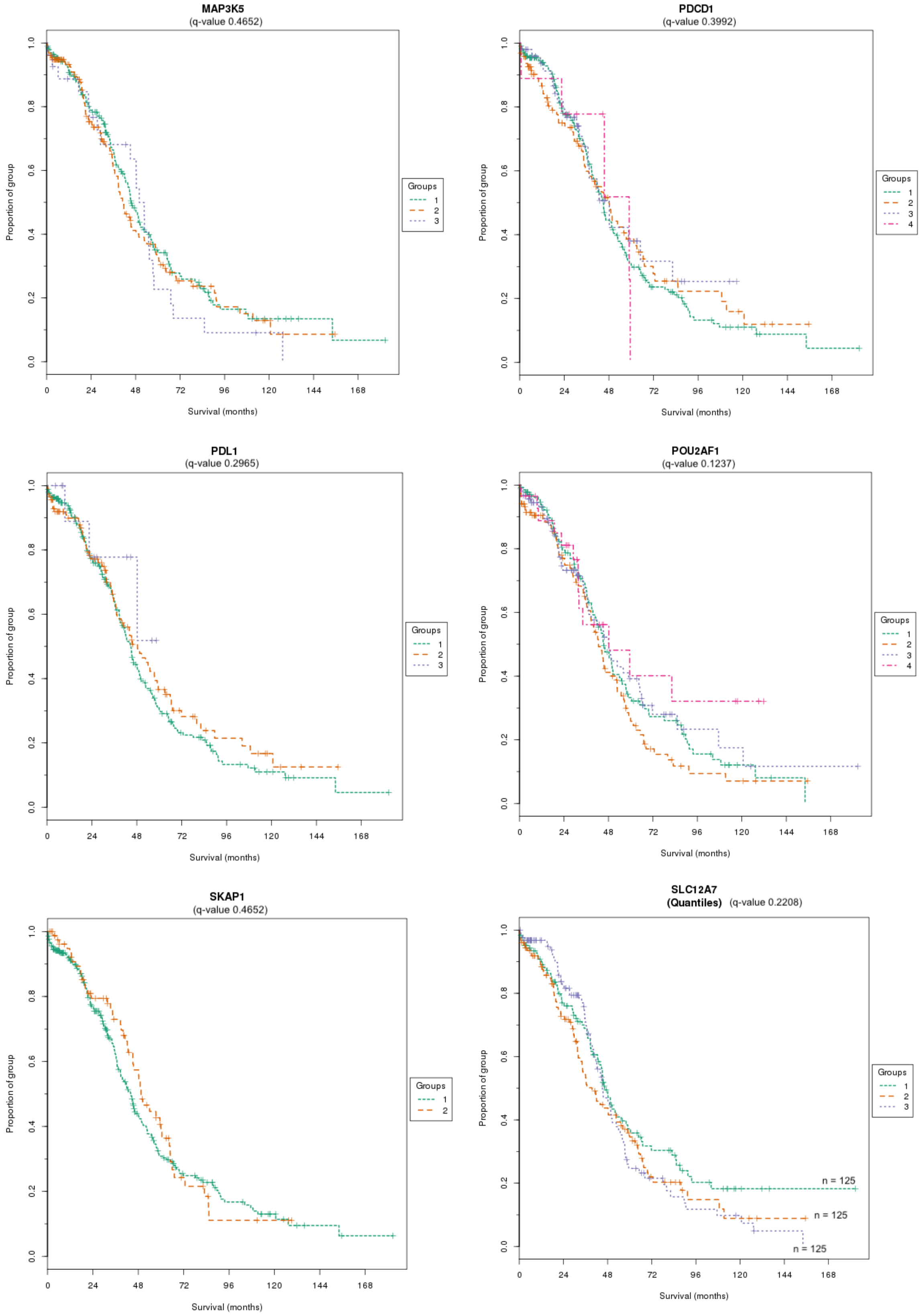


**Figure S5. Survival analysis of BIO_27 in ovarian cancer.** Continued on the next page.

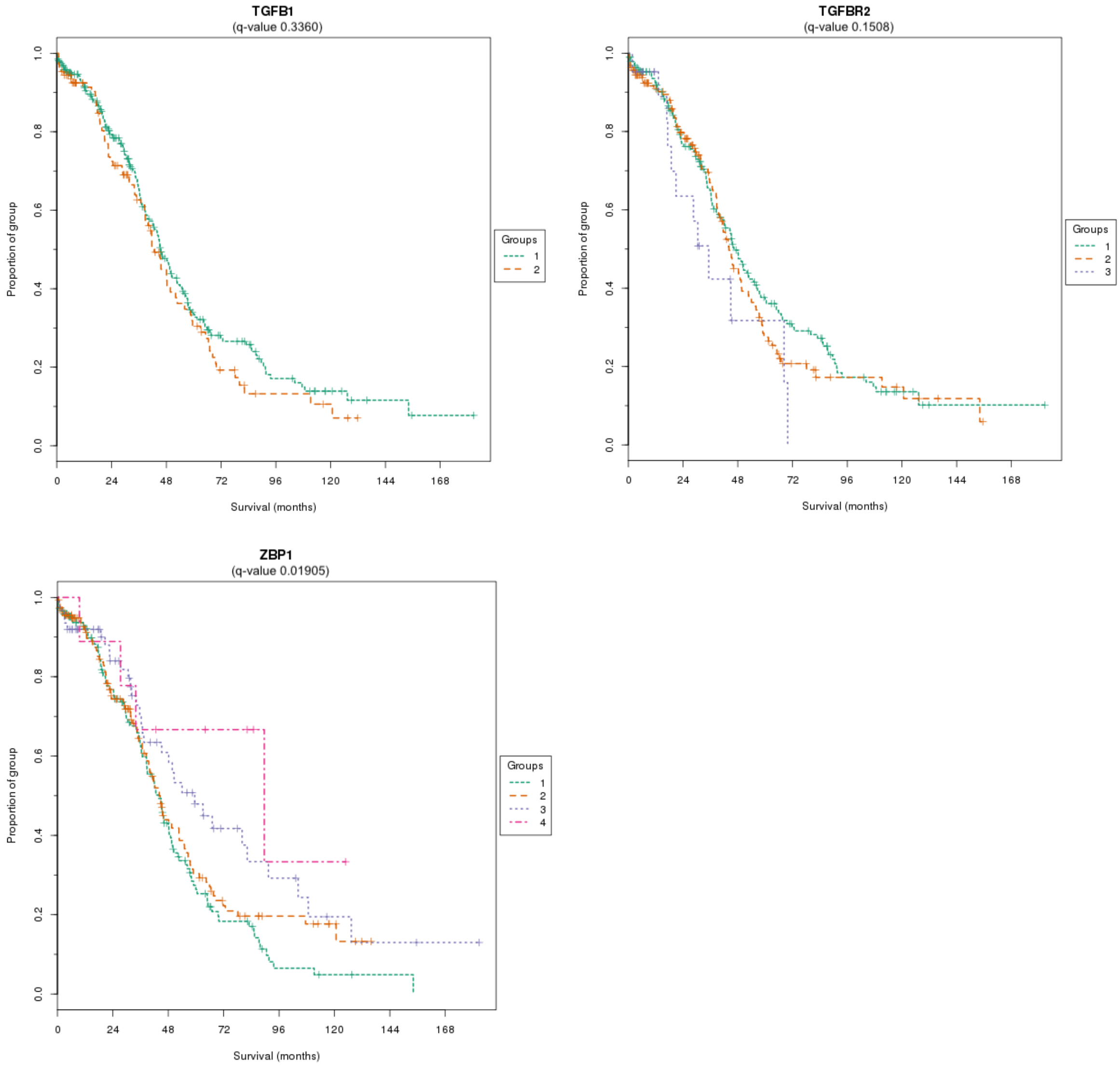


**Figure S5. Survival analysis of BIO_27 in ovarian cancer.** Groups were defined by Gaussian Mixture Modeling of genes expression values in ovarian carcinoma patients from TCGA (n = 375). Numbering in the key identifies each sub-group; higher numbering corresponds to a higher expression level of the gene analysed. Log-rank q-values are given for each gene. A unimodal Gaussian mixture model was returned for SLC12A7, therefore groups were defined using tertiles for this gene. Also see Table 3 in the main manuscript.

| Prognostic factor | Chi-square test | p-value |
|---|---|---|
| | Cutaneous melanoma (n=390) | |
| Age | 1.5323 | 0.22 |
| Tumor stage | 1.5300 | 0.22 |
| CD72 | 0.0382 | 0.85 |
| Overall | 3.5199 | 0.32 |
| | Ovarian carcinoma (n=375) | |
| Age | 11.03 | 0.0009 |
| CD79A | 1.73 | 0.1889 |
| Overall | 13.03 | |

**Table S3. Grambsch–Therneau tests of the proportional hazards assumption.** Results correspond to the multivariate models shown in the main manuscript Tables 4 and 5.
